# Correlates of resting and exertional dyspnea among older adults with obstructive lung disease: a cross-sectional analysis of the Canadian Longitudinal Study on Aging

**DOI:** 10.1101/596114

**Authors:** Joshua Good, Michael K Stickland, Shilpa Dogra

## Abstract

**Introduction:** In patients with chronic obstructive pulmonary disease (COPD), the perception of dyspnea is related to quality of life, and is a better predictor of mortality than the severity of airway obstruction. The purpose of the current study was to use population-level data from the Canadian Longitudinal Study on Aging (CLSA) to identify potential correlates of dyspnea in adults with obstructive lung disease.

**Methods:** Data from participants with a self-reported obstructive lung disease (asthma or COPD) were used for analysis (n=2,854). Four outcome variables were assessed: self-reported dyspnea at 1) rest, 2) walking on a flat surface, 3) walking uphill/climbing stairs, 4) following strenuous activity. Potential sociodemographic, health, and health behaviour correlates were entered in to logistic regression models.

**Results:** Higher body fat percentage, and worse forced expiratory volume in one second were associated with higher odds of reporting dyspnea. Females with an anxiety disorder (OR=1.91, CI: 1.29, 2.83) and males with a mood disorder (OR=2.67, CI: 1.53, 4.68) reported higher odds of experiencing dyspnea walking on a flat surface, independent of lung function and other correlates. Dyspnea while walking uphill/climbing stairs was associated with a slower timed up and go time in females (e.g. OR=1.18, CI: 1.10) and males (OR=1.19, CI: 1.09, 1.30).

**Conclusions:** In addition to traditional predictors such as lung function and body composition, we found that anxiety and mood disorders, as well as functional fitness were correlates of dyspnea. Further research is needed to understand whether targeting these correlates leads to improvements in perceptions of dyspnea.

## INTRODUCTION

Dyspnea, or perceived breathlessness, is a symptom commonly reported by individuals with obstructive lung diseases such as asthma and chronic obstructive lung disease (COPD) ^1, 2^. The prevalence of dyspnea ranges from 9% in the general population ^3^ to approximately 50% in individuals with COPD ^4-7^. These estimates may be low, as dyspnea is often underreported or not recognized by clinicians ^8^. Dyspnea is related to a number of outcomes, including general health ^9^, health related quality of life ^10, 11^, hospital admissions, and mortality ^12^ among individuals with COPD and asthma. In fact in patients with COPD, dyspnea has been shown to be a better predictor of mortality than airway obstruction as measured by the percentage of predicted forced expiratory volume in one second (FEV_1_) ^13^. In other words, the severity of breathlessness cannot simply be predicted by disease severity or lung function measures. Research indicates that the improvement in dyspnea from participation in pulmonary rehabilitation occurs despite a lack of concurrent improvements in FEV_1_ ^14^. This association between dyspnea and physical inactivity is well-established, such that lower physical activity levels are related to worse dyspnea ^15, 16^.

Several other factors may influence perceptions of dyspnea. Recently, Ekström et al. found that female sex, smoking, presence of asthma, and chronic bronchitis were correlates of dyspnea in healthy adults ^3^. FEV_1_, body mass index, anxiety, and depression have also been identified as correlates of resting dyspnea in adults with asthma or COPD ^17-19^. Psychological factors such as mood or anxiety disorders may have particular significance given their increased prevalence among adults with COPD ^20^. However, limited work has been done at the population-level to identify potential predictors of resting and exertional dyspnea among individuals with an existing obstructive lung disease. This is critical, as simple interventions targeting dyspnea may have a significant impact on the quality of life of individuals with obstructive lung diseases. Thus, the purpose of this study was to use population-level data from the Canadian Longitudinal Study on Aging (CLSA) to identify potential correlates of resting and exertional dyspnea. Given that perceived dyspnea is known to differ by sex, analyses were conducted separately for males and females.

## METHODS

### Data Source and Participants

The CLSA is a nationally representative, stratified, random sample of 51,338 Canadian females and males aged 45 to 85 years (at baseline). The purpose of this survey is to collect data on the health and quality of life of Canadians to better understand the processes and dimensions of aging. Data for the present study are from participants (n=30,097) in the Comprehensive sample which collected data through questionnaires, physical examinations, and biological samples.

Inclusion in the CLSA was limited to those who were able to read and speak either French or English. Residents in the three territories and some remote regions, persons living on federal First Nations reserves and other First Nations settlements in the provinces, and full-time members of the Canadian Armed Forces were excluded. Individuals living in long-term care institutions (i.e., those providing 24-hour nursing care) were excluded at baseline; however, those living in households and transitional housing arrangements (e.g., seniors’ residences, in which only minimal care is provided) were included. Finally, those with a cognitive impairment at the time of recruitment were excluded.

Only those with complete data for dyspnea outcomes (n=29,670), sociodemographic variables, selected health conditions, and physical activity (n=18,232) were included in the analysis. Only those who had self reported asthma and/or COPD, and did not report lung cancer were included (n=2,854). Of these, 2,064 had asthma only, 514 had COPD only, and 276 had both asthma and COPD.

Self reported COPD and asthma were determined based on the questions “Has a doctor told you that you have/had any of the following: emphysema, chronic bronchitis, chronic obstructive pulmonary disease (COPD), or chronic changes in lungs due to smoking?” and “Has a doctor ever told you that you have asthma?”. Self report was used rather than spirometry to classify COPD because 24% of participants with self-reported COPD had taken short and/or long acting bronchodilators prior to spirometry.

### Measures

#### Dyspnea Variables

Participants were asked the following four questions relating to resting and exertional dyspnea: “Do you become short of breath walking on flat surfaces?”, “Do you become short of breath climbing stairs or walking up a small hill?”, “Have you had an attack of shortness of breath that came on following strenuous activity at any time within the last 12 months?”, and “Have you had an attack of shortness of breath that came on during the day when you were at rest at any time within the last 12 months?”. Response options to each question were “Yes” or “No”. Each dichotomous variable was treated as an outcome in our analyses.

#### Health and Health Behaviour Variables

##### Chronic Conditions

Participants were asked to report chronic conditions that had been diagnosed by a physician and that were expected to last, or had already lasted, 6 months or more. We chose to use variables pertaining to *anxiety disorder, mood disorder, and cardiovascular disease* due to their known associations with breathlessness ^18, 19, 21^. *Anxiety disorder and mood disorder* were based on the questions “Has a doctor ever told you that you have an anxiety disorder such as a phobia, obsessive-compulsive disorder or a panic disorder?” and “Has a doctor ever told you that you have a mood disorder such as depression (including manic depression), bipolar disorder, mania, or dysthymia?”. Participants were classified as having *cardiovascular disease* if they responded positively to either of the following two questions: “Has a doctor ever told you that you have heart disease (including congestive heart failure, or CHF)?” or “Has a doctor ever told you that you have peripheral vascular disease or poor circulation in your limbs?”.

##### Physical Activity and Functional Fitness

Participants completed a *timed up and go* (TUG) assessment which required them to stand up from a chair with arm rests, walk 3 metres, turn around, walk back, and sit down. The time for participants to complete the TUG was used as a global indicator of physical function. To better understand the impact of exertional dyspnea, we also used responses from the question “In the past 12 months, have you felt like you wanted to participate more in physical activities? If yes, what prevented you from doing physical activities/more physical activities?”. Three response options were selected for the current analysis: “*Injury/Illness*”, “*Health condition limitation*”, and “*Lack of energy*” due to their applicability to individuals with obstructive lung conditions.

##### Others

For *smoking history*, participants were categorized as Never smoked, <10 pack years, or 10 packs years or more based on responses to questions pertaining to the number of cigarettes smoked per day and total years smoked (further detail provided elsewhere ^22^).

*Height* was measured to the nearest 0.1 cm by trained professionals (Seca 213). *Body fat percentage* was measured using bioimpedance (Hologic Discovery A Dual Energy X-Ray Absorptiometry). Total body fat percentage was assessed with participants lying flat on their back with their arms at their side and feet pointed in.

A handheld spirometer was used (TruFlow Easy-On) to assess lung function. Only those with major contraindications did not perform the test ^23^. Maximal inspiratory and expiratory maneuvers were performed to obtain FEV_1_. Further detail on the procedures can be found in the CLSA spirometry standard operating procedures ^23^. Only those who were able to complete three acceptable maximal maneuvers were included; that is, the difference between the best two FEV_1_ and Forced vital capacity values was within 150ml. Participants with extreme data outside of normal physiological limits were also excluded (i.e. FEV_1_ >10 Litres). The best *FEV_1_* was used for analysis. Percent predicted FEV_1_ was calculated based on age, height, ethnicity, and sex, using formulas developed by the Global Lung Function Initiative ^24^. Absolute FEV_1_ was used in analyses rather than percent predicted FEV_1_ due to previous research indicating that absolute values of lung function explain differences in breathlessness ^25^, and because age, sex, and height were already included in each logistic regression.

##### Sociodemographic Variables

Participants were asked to report their *age* and *sex*, and provided information on household income. For *household income*, responses were categorized as “Less than $20,000”, “$20,000 to $49,999”, “$50,000 to $99,999”, “$100,000 to $149,999”, and “$150,000 or more”.

### Statistical Analysis

Means and frequencies were used to describe the sample. Odds ratios were calculated for each of the dyspnea outcomes (at rest, walking on a flat surface, climbing stairs or walking uphill, and following strenuous activity) using logistic regressions. Each of the health and health behaviour (anxiety disorder, mood disorder, heart disease, TUG time, injury/illness limiting PA, health condition limiting PA, lack of energy limiting PA, smoking history, height, body fat percentage, and FEV_1_), and sociodemographic (age and household income) variables were included in the same models. Analyses were conducted separately for males and females ^3^.

All analyses were performed using SPSS v.24. To ensure national representation and to compensate for under-represented groups, sampling weights were applied to regression models. Significance was set at p<0.05. Additional details on sampling, methods and weighting on the CLSA can be found in the protocol document ^26, 27^.

## RESULTS

The sample had a mean age of 61.3 ± 9.8 years and was 60.0% female. Females had a significantly higher body fat percentage, were shorter, had a lower FEV_1_, and a higher percent predicted FEV_1_ (Table 1). More females also reported shortness of breath at rest, walking on flat surface, and walking uphill/climbing stairs than males. Additional sample characteristics can be found in Table 1.

**Table 1:** Sample characteristics of adults with obstructive lung disease

| Characteristics |  | Males (n=1,139) | Females (n=1,715) |
| --- | --- | --- | --- |
| Age (years) | N/A | 61.4 ± 9.9 | 61.2 ± 9.7 |
| Body fat percentage | N/A | 28.8 ± 5.8 | 40.2 ± 6.4* |
| Height (cm) | N/A | 175.0 ± 6.8 | 161.6 ± 6.6* |
| FEV<sub>1</sub> (L) | N/A | 3.0 ± 0.7 | 2.2 ± 0.6* |
| Percent predicted<br>FEV<sub>1</sub> (%) | N/A | 86.9 ± 17.3 | 89.0 ± 16.6* |
| FEV<sub>1</sub>/FVC | N/A | 73.1 ± 8.3 | 74.8 ± 7.0* |
| Percent below lower<br>limit of normal for<br>FEV<sub>1</sub>/FVC | N/A | 14.8% | 9.3% |
| Household income<br>(% of sample)^ | Less than \$20,000 | 3.9% | 7.1% |
| | \$20,000 to \$49,999 | 16.0% | 26.1% |
| | \$50,000 to \$99,999 | 33.5% | 34.9% |
| | \$100,000 to \$149,999 | 24.3% | 16.6% |
| | \$150,000 or more | 22.3% | 15.4% |
| Smoking history | Never smoked | 41.6% | 43.9% |
|  | <10 Pack Years | 23.8% | 25.5% |
|  | 10 or more Pack Years | 34.6% | 30.6% |
| Anxiety disorder (%<br>yes)^ | N/A | 9.1% | 12.9% |
| Mood disorder (%<br>yes)^ | N/A | 17.4% | 26.1% |
| Heart disease (%<br>yes) | N/A | 16.6% | 13.9% |
| Timed up and go<br>(seconds to<br>complete) | N/A | 9.4 ± 2.0 | 9.5 ± 2.6 |
| Factor preventing<br>more participation in<br>physical activity (%<br>yes) | Injury/Illness^ | 9.8% | 12.5% |
|  | Health condition<br>limitation^ | 11.7% | 14.6% |
|  | Lack of energy^ | 4.1% | 6.0% |
| Reported shortness<br>of breath (% yes) | At rest^ | 6.2% | 9.0% |
|  | Walking on a flat<br>surface^ | 9.2% | 16.1% |
|  | Walking uphill/climbing<br>stairs^ | 27.6% | 43.6% |
|  | Following strenuous<br>activity | 27.0% | 29.0% |

Note: Data for continuous variables are presented as mean ± standard deviation. ^p<0.05 for the chi-square for categorical variables; \*p<0.05 using t-tests

For all logistical regressions, OR presented are from models where all variables were entered simultaneously, thus these findings are independent of lung function, height, age, and other correlates.

Significant associations in the sample of males are highlighted in Figure 1. Having a mood disorder was associated with higher odds of reporting dyspnea at rest, while walking on a flat surface, and following strenuous activity when compared to males without a mood disorder. A one litre higher FEV_1_ was associated with approximately half the odds of reporting dyspnea for three of four dyspnea outcomes.

**Figure 1:**
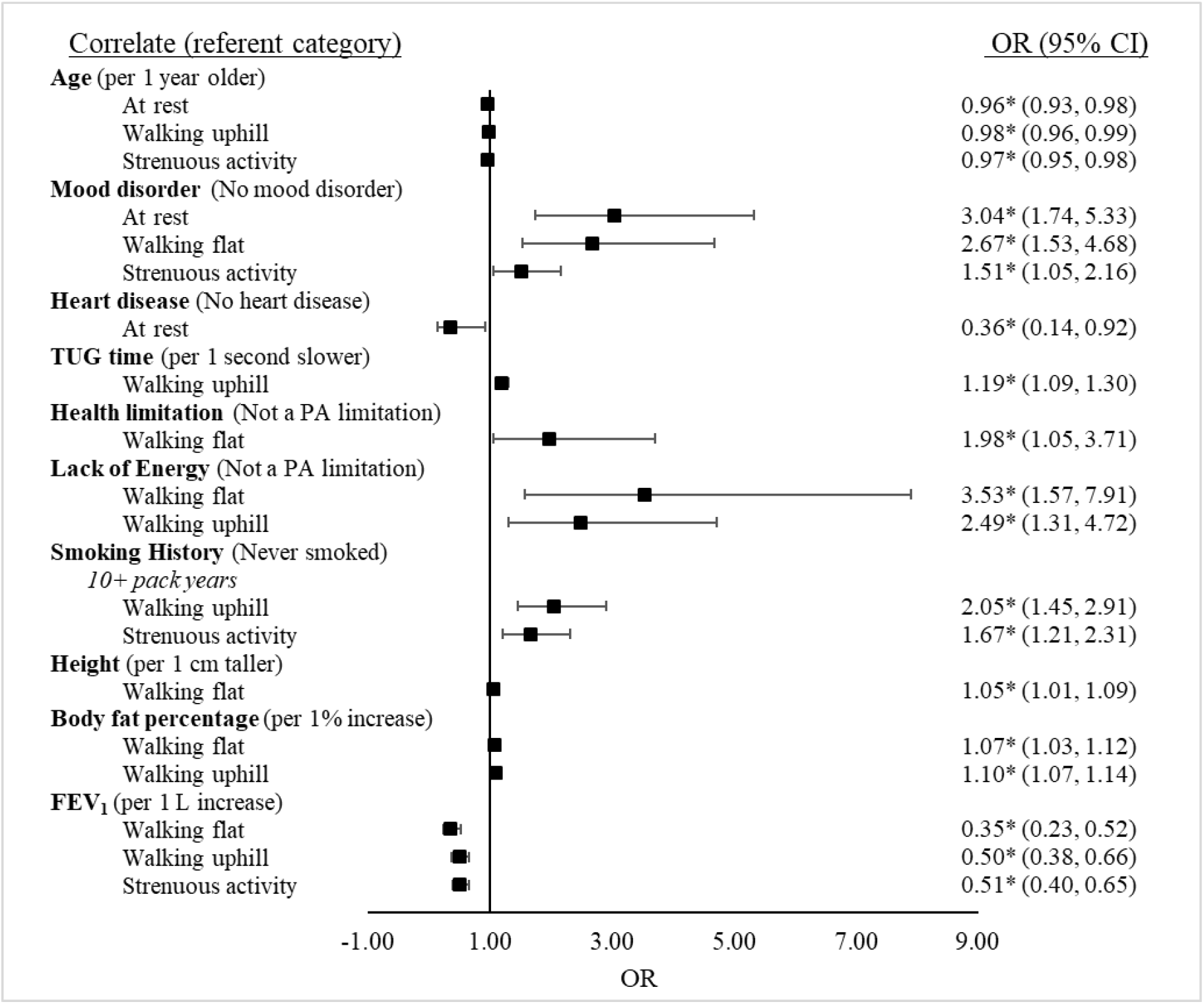
Significant associations of self-reported resting and exertional dyspnea in the past year with sociodemographic, health and health behavioural predictors among males with obstructive lung disease (n=1,139) Note: A higher OR indicates a higher odds of reporting dyspnea at rest, walking on a flat surface, walking uphill/climing stairs, or following strenuous activity relative to the referent category. *p<0.05

Significant associations in the sample of females are highlighted in Figure 2. Females with an anxiety disorder were more likely to report dyspnea while walking on a flat surface, and following strenuous exercise compared to those without an anxiety disorder. A slower TUG time was also associated with higher odds of reporting dyspnea for all three exertional dyspnea outcomes.

**Figure 2:**
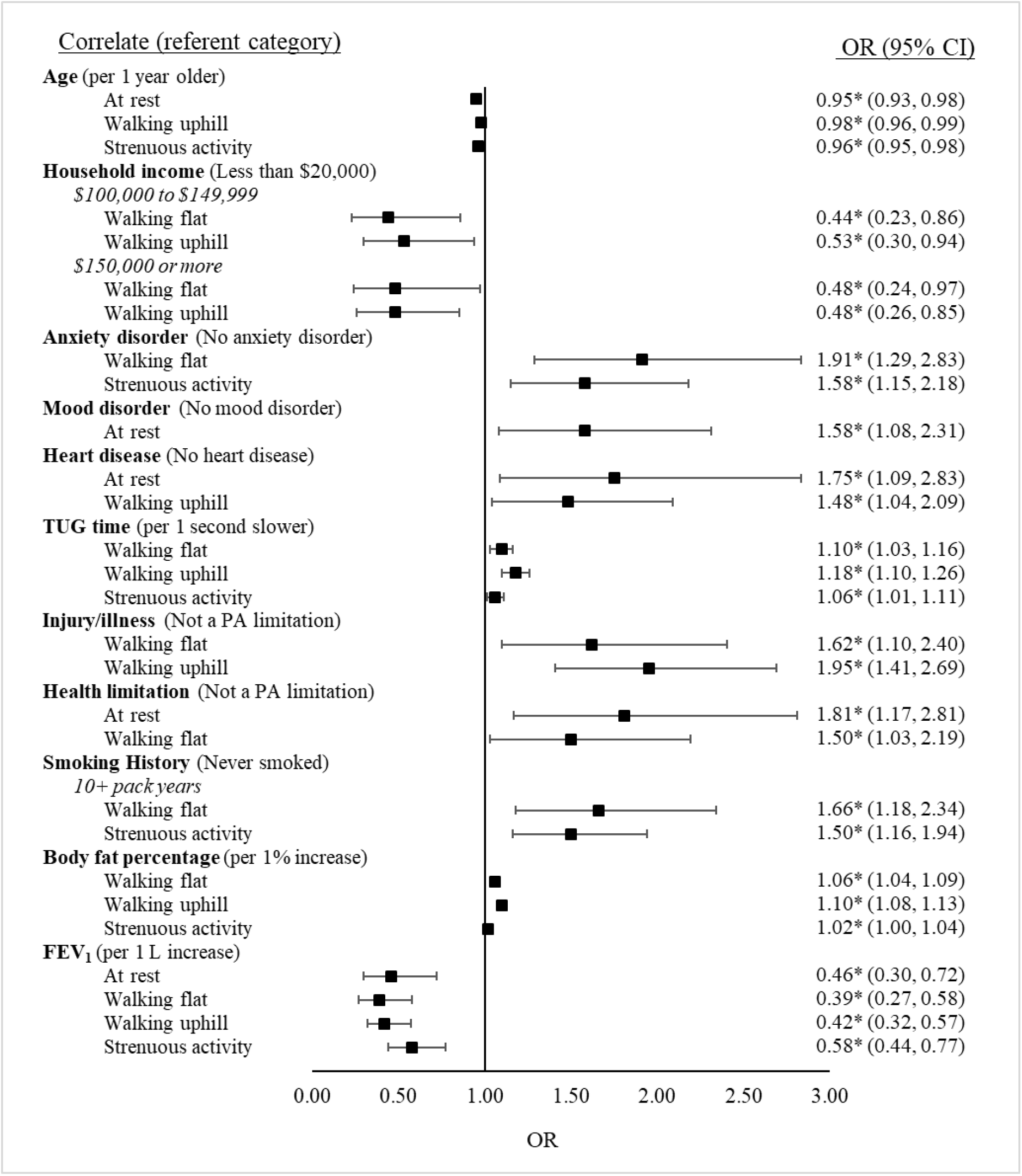
Significant associations of self-reported resting and exertional dyspnea in the past year with sociodemographic, health and health behavioural predictors among females with obstructive lung disease (n=1,715) Note: A higher OR indicates a higher odds of reporting dyspnea at rest, walking on a flat surface, walking uphill/climbing stairs, or following strenuous activity relative to the referent category. *p<0.05

## DISCUSSION

Using data from the CLSA, we sought to identify correlates of resting and exertional dyspnea among middle-aged and older Canadian males and females with an obstructive lung disease. The three main findings of this study are 1) females with an anxiety disorder and males with a mood disorder have higher odds of reporting dyspnea, independent of lung function and other correlates, 2) functional fitness may be important for limiting dyspnea, as individuals, particularly females, with a slower TUG time were more likely to report dyspnea, 3) age, body fat percentage, and FEV_1_ are important correlates of dyspnea among both males and females. These findings are the first to use population level data to identify potential correlates of dyspnea in adults with asthma and COPD, and have implications for future research and interventions targeting dyspnea.

### Health and Health Behaviours

Chronic Conditions: With regards to the chronic condition variables assessed, we found that having a mood disorder, anxiety disorder, or cardiovascular disease was associated with a higher odds of reporting dyspnea, even after adjusting for the effects of lung function, body fat %, and other correlates.

Previous research on the association between dyspnea and anxiety or depressive symptoms is mixed among individuals with COPD. Most studies demonstrate an association ^18, 28, 29^; however, some do not ^30^. It has been suggested that the stress of having a chronic condition may be related to the increased likelihood of having an anxiety or mood disorder ^31^. For anxiety disorders in particular, the hyperventilation model has been suggested, whereby anxiety results in baseline hyperventilation, which when added to obstruction and hypocapnia can aggravate symptoms of dyspnea ^32^. Patients with anxiety may also be more likely to misinterpret symptoms of mild dyspnea as more severe ^32^. Further, highly anxiety sensitive individuals appear to be more sensitive to the anticipation of a hyperventilation procedure ^33^. Therefore, the anticipation of dyspnea may lead to a greater perception of dyspnea among individuals with both anxiety and an obstructive lung disease.

We observed sex-differences in the associations with anxiety and mood disorders. The reason for these sex differences is unclear. It has been suggested that males and females with major depression present with different symptoms ^34^. Females also have higher rates of both depression and anxiety ^35^. The differences observed in the current study may relate to the subjective experience of how males and females experience dyspnea. Males with mood disorders may be more likely to experience dyspnea, whereas females may experience different symptoms. In our study, the prevalence of anxiety was similar between males and females, but mood disorders were more prevalent in females compared to males (26 vs. 17%). This may help to explain sex differences observed in mood disorder. Understanding these associations is critical, as depression and anxiety are often untreated in adults with COPD ^36^. Thus, proper treatment of anxiety and mood disorders may be aid in improving sensations of dyspnea, and thus quality of life.

The presence of heart disease was associated with dyspnea in females, such that having heart disease was associated with higher odds of reporting dyspnea at rest and while walking uphill ^3^. However, among males, only dyspnea at rest was associated with heart disease, and this association was in the opposite direction expected. It is possible that the variable we used was not appropriate as it included any heart disease, including congestive heart failure. Future research using specific cardiovascular disease variables may help us understand the interactive effects of cardiopulmonary disease on dyspnea.

Physical Activity and Functional Fitness: Reporting factors (Injury/Illness, Health condition limitation, and Lack of energy) that limited participation in physical activity, or having a slower TUG time, were both associated with reporting dyspnea. Among males, reporting a lack of energy that prevented more participation in physical activity was associated with dyspnea while walking on a flat surface and while walking uphill/climbing stairs. This is not surprising, as fatigue, or lack or energy, is a common symptom among those with COPD, and has been shown to be correlated with dyspnea, ^37-39^.

Among females, TUG time, injury/illness, and reporting a health limitation that prevented more participation in physical activity were the most consistently associated with resting and exertional dyspnea. TUG time may be an important modifiable correlate of dyspnea as the presence of dyspnea may lead to avoidance of physical activity, causing further deconditioning, and thus worsening of dyspnea. Self-reported functional fitness has been shown to be associated with dyspnea in adults with COPD ^37, 40 41^. Pulmonary rehabilitation programs have also led to improvements in 6 minute walk distance, and dyspnea ^14, 42, 43^. Thus, measures of functional fitness may be useful clinical tools for physicians to use to better understand dyspnea in their patients.

Others: Our finding that body fat percentage was associated with exertional dyspnea outcomes is consistent with previous literature that indicates that fat free mass index is correlated with dyspnea among COPD patients ^44^, and that body mass index has is associated with dyspnea among adults 40 years and older ^3, 4, 45^. It is important to note that higher levels of exertional dyspnea among individuals with more body fat may be related to reduced lung volumes and greater work of breathing required ^45^; however, body mass index is often lower in individuals with the most severe COPD, (GOLD stages 3 and 4) ^46^. Clearly, body fat is not the only factor leading to an increased perception of dyspnea, but it may be a contributing factor.

An interesting sex-difference emerged on the association of FEV_1_ and dyspnea outcomes. Dyspnea at rest was significantly associated with FEV_1_ among females, but not among males. Females may be more prone to reporting dyspnea at rest than males due to smaller airways, and smaller lung volumes ^47, 48^. In fact, in our sample, only 71 of 1,139 males (6.2%) reported dyspnea at rest. For all exertional dyspnea outcomes, a lower FEV_1_ was significantly associated with a higher odds of reporting dyspnea (after adjustment for age and height). FEV_1_ has consistently been shown to be negatively associated with dyspnea among the general population ^3, 4^, individuals with asthma ^11^, and with severe COPD ^49^. Despite these associations, the relevance of this is unclear as even in the absence of lung function improvements following pulmonary rehabilitation, improvements in dyspnea have been observed ^14^.

### Sociodemographic Correlates

Our finding that older age was associated with lower dyspnea was of interest. Increasing age is associated with a decline in lung function ^24^ and some studies from the general population indicate that increasing age is associated with an increase in severity or prevalence of dyspnea ^6, 7^. However, there is evidence to suggest that increasing age is associated with a decrease in the occurrence of dyspnea ^4^. This supports the notion that individuals can become accustomed, and therefore desensitised to feelings of breathlessness. In fact, researchers have suggested a decrease in the perception of bronchoconstriction in response to methacholine in older adults with and without asthma ^50^. Furthermore, it is possible that individuals with dyspnea are less likely to survive to old age, leaving an older population that is “healthier” and less likely to report dyspnea (survivor bias) ^6^. This emphasizes the importance of treating dyspnea seriously in individuals of all ages.

Our finding that household income was only significant for two of the dyspnea outcomes among females, and with none of the outcomes among males was somewhat surprising. Previous research indicates that higher household income and lower social disadvantage is associated with lower perceived dyspnea ^45, 51^. However, these studies assessed dyspnea using the mMRC, and Bowden et al. used social disadvantage based on geographical area rather than household income. Importantly, socioeconomic status of the sample used for the current study was high, with 72.1% of participants reporting a household income greater than $50,000. Thus, future research in a more diverse sample may be warranted.

### Strengths and Limitations

Strengths of this study include the large representative sample, and the objectively measured lung function and body composition measures. There are also a number of limitation to the present study. First, the questions related to dyspnea for the CLSA have not been used in other research studies so direct comparisons can not be performed. The CLSA has performed a validation study for the broader set of chronic airflow obstruction questions and spirometry, regarding the ability to detect asthma or COPD ^52^. Second, while we did remove those with lung cancer, we did not adjust for additional chronic conditions. Thus, care should be taken when interpreting the findings, as other chronic conditions may impact perceptions of dyspnea. Third, participants in the present study had fairly mild obstruction (86.9 ± 17.3 and 89.0 ± 16.6 percent of predicted FEV_1_ for males and females, respectively), possibly due to the inclusion of individuals who may have well controlled asthma. However, this emphasizes that the perception of dyspnea in this sample was not only due to poor lung function. Finally, it is important to note that data from the CLSA are cross-sectional, thus, reverse-causality cannot be ruled out at this time.

## CONCLUSION

In conclusion, using data from a large Canadian sample of older adults with obstructive lung disease, we found that those with an anxiety or mood disorder, and those with lower functional fitness had higher odds of reporting dyspnea after adjusting for lung function and other important correlates. These findings have implications for clinical practice as they could aid in early identification of adults at increased risk of dyspnea and poor health outcomes. This study also provides a basis for future research exploring the role of functional fitness and physical activity in improving perceptions of dyspnea among those with existing obstructive lung disease.

## Acknowledgements

The opinions expressed in this manuscript are the author’s own and do not reflect the views of the Canadian Longitudinal Study on Aging.

This research was made possible using the data collected by the Canadian Longitudinal Study on Aging (CLSA). Funding for the CLSA is provided by the Government of Canada through the Canadian Institutes of Health Research (CIHR) under grant reference: LSA 9447 and the Canada Foundation for Innovation. This research has been conducted using the CLSA dataset Baseline Comprehensive version 3.1 and Maintaining Contact version MCQ v2.0, under Application Number 170315. The CLSA is led by Drs. Parminder Raina, Christina Wolfson, and Susan Kirkland.

This work was supported by the Canadian Institutes of Health Research [funding reference number 372547]. The funding body was not involved in the design of the study, collection, analysis, interpretation of data, and in writing the manuscript.

## Conflict of interests

None declared

